# Highly multiplexed quantitative phosphosite assay for biology and preclinical studies

**DOI:** 10.1101/2020.12.08.415281

**Authors:** Hasmik Keshishian, E. Robert McDonald, Randy Melanson, Dale A. Porter, Filip Mundt, Karsten Krug, Luke Wallace, Dominique Forestier, Bokang Rabasha, Sara E Marlow, Judit Jane-Valbuena, Ellen Todres, Harrison Specht, Javad Golji, Eric Kuhn, Michael Burgess, Melanie MacMullan, Shankha Satpathy, D.R. Mani, Michael Gillette, Karl Clauser, Tomas Rejtar, Karen Wang, Levi A. Garraway, William R. Sellers, Steven A. Carr

## Abstract

Reliable methods to quantify dynamic signaling changes across diverse pathways are needed to better understand the effects of disease and drug-treatment in cells and tissues but are presently lacking. Here we present SigPath, a targeted mass spectrometry (MS) assay that measures 284 phosphosites in 200 phosphoproteins of biological interest. SigPath probes a broad swath of signaling biology with high throughput and quantitative precision. We applied the assay to investigate changes in phospho-signaling in drug-treated cancer cell lines, breast cancer preclinical models and human medulloblastoma tumors. In addition to validating previous findings, SigPath detected and quantified a large number of differentially regulated phosphosites newly associated with disease models and human tumors at baseline or with drug perturbation. Our results highlight the potential of SigPath to monitor phosphoproteomic signaling events and to nominate mechanistic hypotheses regarding oncogenesis, response and resistance to therapy.

## Introduction

Cellular processes including signal transduction, cell cycle progression, and response to DNA damage among many others are regulated through the addition or removal of phosphate from the amino acids serine, threonine and tyrosine. In keeping with this, aberrant phospho-signaling is a hallmark of many diseases including cancer. For example, genetic disruption of the tumor suppressors PTEN and APC leads to pathologic levels of phosphorylated AKT and ß-catenin respectively, while oncogenic activation of ABL and RAS lead to aberrant phosphorylation in CRKL or MEK, respectively. Dysregulated kinases and phosphatases have thus become important targets for therapeutic development. This in turn has motivated the desire to quantitatively monitor phosphorylation events to determine the cellular or organismal activity of such inhibitors. Unfortunately, due to the limited ability to robustly quantify hundreds of phosphorylation events, most drug discovery programs in this area have followed single phosphorylation events as the marker of pharmacodynamic activity. As a result, paradoxical activation of RAF isoforms as a consequence of BRAF inhibitors (Hatzivassiliou et al., 2010; Poulikakos et al., 2010), or feedback upstream pathway activation resulting from mTOR inhibitors were missed until well after the relevant molecules were in clinical trials or beyond (Shi et al., 2005).

The Cancer Cell Line Encyclopedia project (CCLE), in addition to characterizing genome, transcriptome and methylome alterations (Barretina et al., 2012; Ghandi et al., 2019) has sought to characterize the metabolome and proteome across hundreds of cancer cell lines (Li et al., 2019; Nusinow et al., 2020). An initial attempt in the phosphoproteome space was also made using Reverse Phased Protein Arrays (Li et al., 2017), however, the sparse availability of phosphoantibodies that are highly reliable in detecting phosphorylation events on the protein arrays limits the broader application of this approach. Thus, to begin to develop high complexity quantitative phosphoprotein assays for use in both characterizing cell lines and clinical samples, and to enable much deeper pharmacodynamic assessment of therapeutics, we set out to develop robust mass spectrometry based phospho-assay sets.

Mass spectrometry (MS)-based proteomics has led to the discovery of the majority of the over 200,000 known human phosphosites (Phosphosite.org (Hornbeck et al., 2015)). Many laboratories, but especially the Clinical Proteomics Tumor Analysis Consortium ([CPTAC](https://proteomics.cancer.gov/programs/cptac)), have elaborated deep, high quality phosphopeptide and proteome datasets (Archer et al., 2018; Chen et al., 2017; Dou et al., 2020; Gillette et al., 2020; Huang et al., 2017; Mertins et al., 2016; Satpathy et al., 2020; Vasaikar et al., 2019; Zhang et al., 2014; Zhang et al., 2016). The deepscale proteomic methods used in the CPTAC studies detect 30,000-45,000 distinct phosphosites in each sample studied, and quantitative chemical labeling provides relative quantification of each site across samples. However, one drawback of this approach is the lack of uniform detection of any given phosphosite across an entire sample cohort, a technical artifact caused by stochastic sampling of the analytes introduced into the MS system, especially in low abundance, and the extreme complexity of the samples analyzed.

Targeted MS in the forms of multiple reaction monitoring (MRM, also referred to as selected reaction monitoring, SRM) and parallel reaction monitoring (PRM) are now widely-used for highly multiplexed, quantitative measurement of proteins in blood, cells and tissues (Abelin et al., 2016; Addona et al., 2011; Eshghi et al., 2020; Gallien et al., 2015; Huttenhain et al., 2019; Keshishian et al., 2009; Kuhn et al., 2009; Sperling et al., 2019; Whiteaker et al., 2018). All variants of the approach begin with targeted selection in the mass spectrometer of the intact, ionized peptides of interest followed by fragmentation of each peptide precursor to produce product ions that, together with the mass of the intact peptide, are used to identify and quantify that peptide. In its most specific and precise form, heavy isotope-labeled synthetic peptides are added at known concentration to verify that the correct peptide is being measured and to improve accuracy of the relative quantification of target peptide. The use of this technology to measure posttranslationally-modified peptides is less common. Targeted MS assays have largely been developed for purposes of verification, however, if such an assay queries a large number of targets it can also be viewed as a discovery method.

In contrast to multiplexed antibody methods, MS-based targeted analysis of peptides and modified peptides, including phosphopeptides, can, in principle, be configured to quantify any phosphosite of interest in any organism and scaled to measure many hundreds of peptides in a single measurement cycle of a few hours by LC-MS/MS (Burgess et al., 2014). The methods have been extended in two recent studies to detect and quantify on the order of 100 phosphosites in a single 1-2 h analysis (Abelin et al., 2016; Kennedy et al., 2016). Both of these studies employed immobilized metal affinity to enrich phosphopeptides from cells followed by analysis of the resulting mixture by targeted MS using heavy stable isotope-labeled phosphopeptides for confident detection and quantification. Kennedy et al. measured phosphosites relevant to DNA damage response (Kennedy et al., 2016), while Abelin et al. assayed a set of readily detectable, moderately abundant phosphosites known to be modulated in non-uniform ways by a panel of drugs in a range of cell lines (Abelin et al., 2016).

Here we present the development and application of the largest phosphosite assay to date, which we call SigPath. This quantitative, targeted MS-based assay measures 284 phosphosites in 200 cancer-relevant target proteins spanning many pathways. Phosphosites were selected by cancer biologists based on known or presumed relevance to cancer disease or treatment and through discovery proteomics efforts leading to the SigPath set of 295 phosphopeptides.

This unique set is purposely designed to probe a broad swath of signaling biology in a single measurement rather than a focusing on a single pathway. Importantly, the panel can be extended to measure other phosphosites in additional pathways as desired. Here, we demonstrate the utility of the assay through a range of applications in drug-treated cells lines, preclinical models of breast cancer and human medulloblastoma tumor samples.

## Results

### Selection of phosphosites for assay development and assay construction

The majority of the phosphosites were nominated by cancer biologists in our institutions and supplemented with frequently modulated phosphosites observed in our discovery proteomics experiments (Figure S1). In an effort to further characterize the signaling and physiological state of human tumor samples, we set out to capture nodes of biological pathways known to be modulated via phosphorylation. Critical kinase cascades, often hyperactivated in tumors, make up the backbone of the assay set. These include the MAPK, PI3K, PKC, SRC and JAK signaling pathways as well as both receptor and non-receptor tyrosine kinases. Both critical activation sites of the kinases themselves and their relevant downstream substrates were targeted. These kinase cascades often culminate in the modulation of transcription factor circuits, so transcriptional nodes within the FOXO, STAT, NFKB, TGFB and Wnt pathways were also selected. Lastly, to capture physiological cell states, protein phosphosites were selected to provide readouts on DNA damage, cell cycle arrest, apoptosis, spindle checkpoint activation, hypoxia, autophagy, cell stress and epithelial-to-mesenchymal transition. The rationale for specific target selection and pathway information for the selected sites is presented in Tables S1A and S1B, respectively. While these sites were selected based on relevance to cancer cell-autonomous phenotypes, the associated pathways constitute central signaling nodes and should therefore be valuable when applied to a variety of experimental paradigms.

### SigPath assay development

Assay development, configuration, quality assessment as well as workflow development (Figure S1A, 1A) are described in Star Methods. The overall success rate for detection of the targeted phosphopeptides was 85% (Figure S1B). Twenty-four out of 352 phosphopeptides failed assay configuration due to their LC or MS characteristics, while another 30 failed during the phosphotyrosine (pY) antibody or immobilized metal affinity chromatography (IMAC) enrichment steps (Table S1A). It is noteworthy that of the 37 phosphopeptides that were included in the assay panel but lacked prior experimental observation in our datasets (mostly pY-containing), we were able to successfully configure assays for 24 of them. The final working assay contains 298 phosphopeptides representing 284 phosphosites and 200 phosphoproteins (Table 1 and Table S1A). The SigPath panel predominantly consists of singly-phosphorylated peptides, with only 8 doubly-phosphorylated phosphopeptides included. The majority of phosphoproteins (142/200) are represented by a single tryptic phosphopeptide containing a single phosphosite (Figure S1C).

**Table 1:**
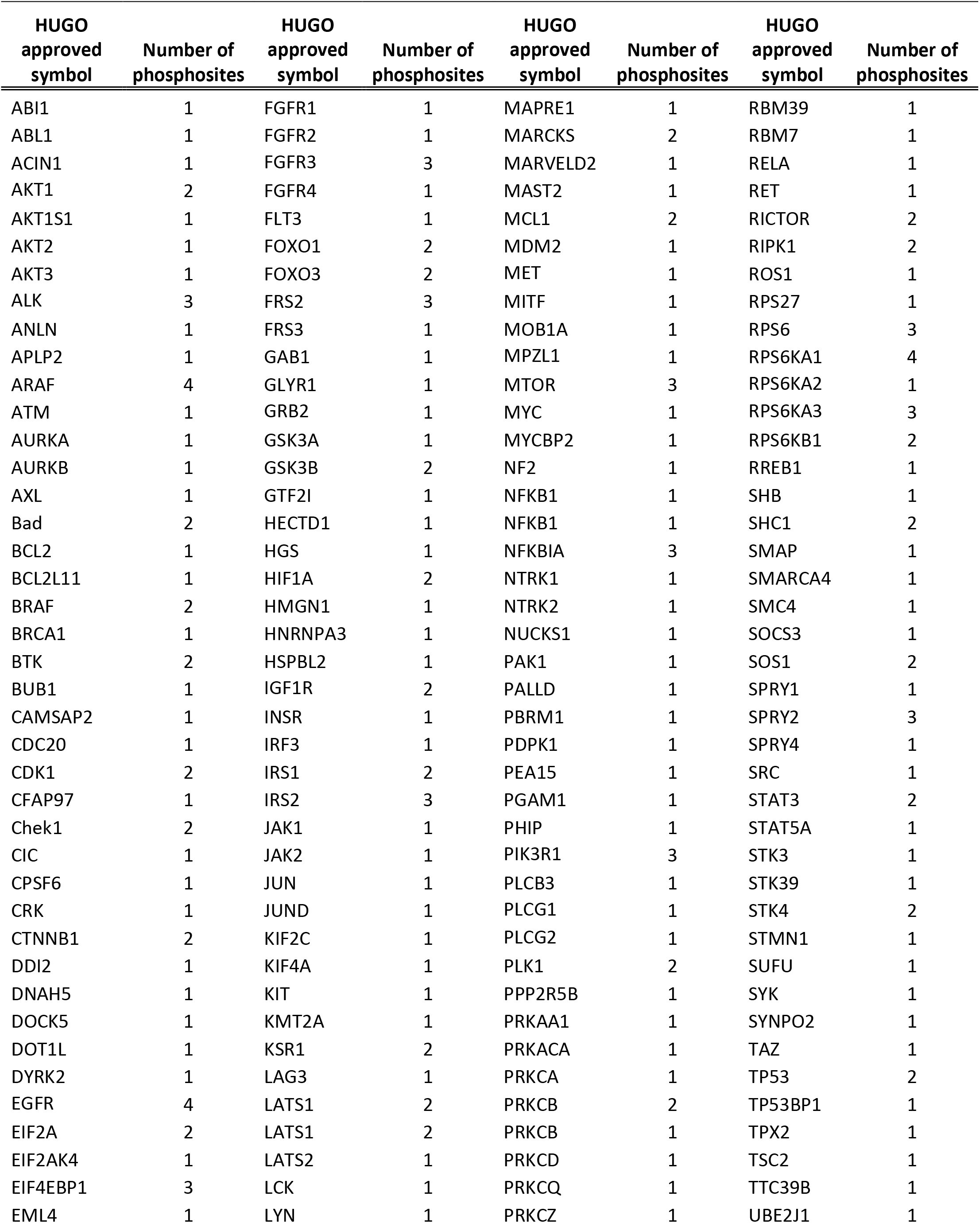
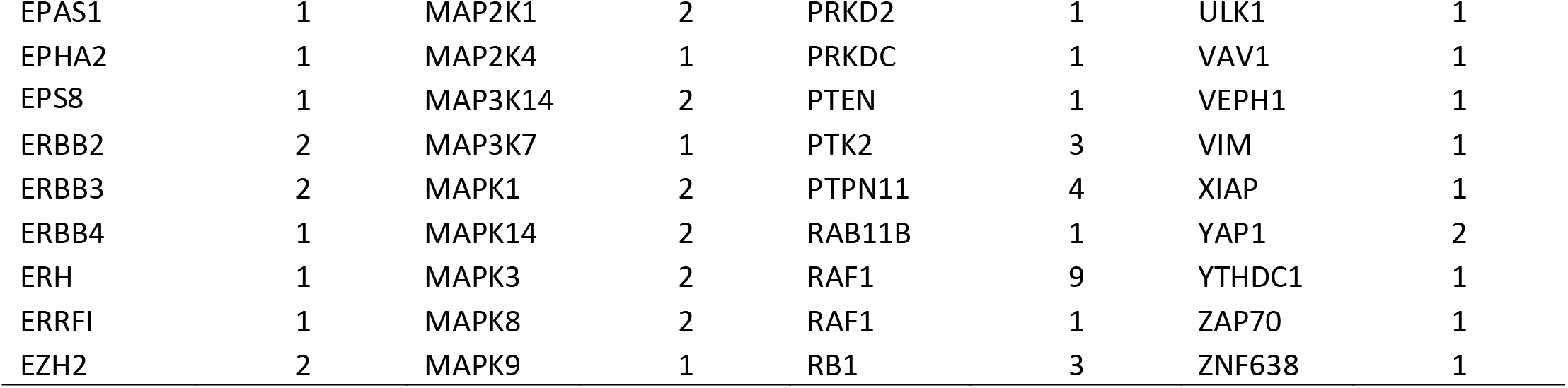
List of all the proteins in the final SigPath assay panel. Table includes HUGO gene names and number of phosphosites per protein in the assay panel.

### Pathway representation in the final assay panel

Pathways represented by the final SigPath panel were assessed using the canonical databases in the Molecular Signatures Database (MSigDB)((Jassal et al., 2020; Kanehisa and Goto, 2000; Liberzon et al., 2015; Liberzon et al., 2011; Pico et al., 2008; Schaefer et al., 2009). The panel represents a spectrum of cancer-relevant biology spanning signal transduction, cell proliferation, apoptosis and the immune system (Table S1B). Figure 1B shows range of pathways represented in Hallmark gene sets and figures 1C and 1D show coverage of EGFR and FGFR pathways, respectively.

**Figure 1.**
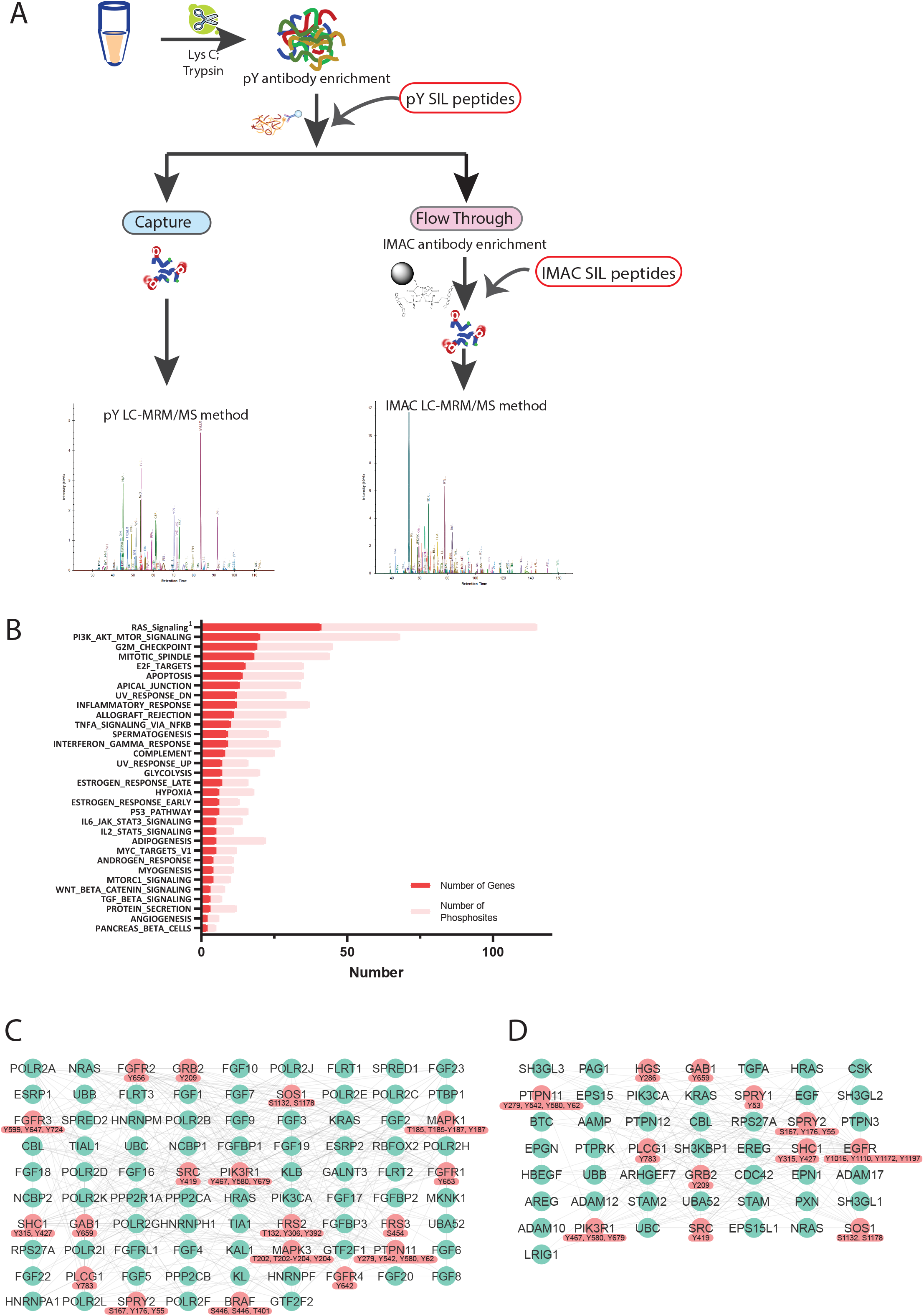
SigPath assay workflow and pathway coverage (A) After digestion of the sample, it’s spiked with pY and IMACpY mixtures of SIL peptides and enriched using pY antibody. Portion of the flow through is spiked with the IMAC mixtures of SIL peptides and enriched by IMAC. Both, pY and IMAC captured samples are analyzed on the MS using pY and IMAC LC-MRM/MS methods, respectively. (B) Pathways represented by SigPath in MSigDB Hallmark pathway category. To be included in the plot a pathway had to have at least 5% coverage, or be represented by a minimum of 3 proteins and 5 phosphosites in the assay. Both, number of genes (red) and phosphosites (pink) are shown on the plot. ^1^ Included from MSigDB WikiPathway pathway category. (C) Signaling by EGFR pathway (MSigDB Reactome) coverage in SigPath assay. Colored in red are all proteins and phosphosites included in the SigPath assay. (D) Signaling by FGFR pathway (MSigDB Reactome) coverage in SigPath assay. Colored in red are all proteins and phosphosites included in the SigPath assay.

### Application of the assay to evaluate effects of drug treatment in cancer cell lines

We initially evaluated the assay in ten cell lines representing various cancer types (lung, B-cell lymphoma, mantle cell lymphoma, prostate, ovarian, bladder and melanoma) and genetic contexts using 250 of the 298 phosphopeptides in SigPath (Table S2A). Each cell line was selected to maximize the potential for detecting endogenous signals from the phosphosites in the panel that are typically at low abundance. Endogenous versions of 81 to 126 peptides were detected in each individual cell line with a total of 160/250 (70%) detected across all 10 cell lines. Detection of endogenous phosphopeptides reflected the genetic context of the cell lines (Figure S2A). For example, phosphosites/peptides derived from ALK and FGFR were only detected in lung adenocarcinoma cell line H3122 (driven by *ALK* fusion) and bladder carcinoma cell line RT112 (driven by *FGFR3* fusion), respectively, while in lung adenocarcinoma cell line, PC9, driven by *EGFR* mutation, higher levels of EGFR phosphosites were detected.

To investigate the utility of the SigPath assay to detect and quantify acute perturbations in cell lines, we next treated lung adenocarcinoma (LUAD) H3122 cells with the ALK inhibitor Ceritinib (Friboulet et al., 2014; Shaw et al., 2014) and KRAS mutant Ls513 colorectal cancer cells with the MEK inhibitor Trametinib (Falchook et al., 2012; Flaherty et al., 2012) (Figure S2B – S2C). Cells were treated with the respective inhibitors or DMSO for 6 and 24 hours and two process replicates per condition were analyzed using the SigPath assay as described above. A total of 182 and 163 endogenous phosphopeptides were detected in H3122 (lung) and Ls513 (colon) cell lines, respectively (Table S2B). Excellent reproducibility was achieved for replicates with Pearson correlation of greater than 0.9 for all samples in both cell lines (Figure S2D).

Ceritinib treatment of H3122 cells resulted in significant regulation of 44 phosphosites, 80% of which were downregulated with 24 hours of drug treatment (Figure 2A). Consistent with ALK inhibition, several pY sites on ALK (Y1096, Y1507, Y1604) showed dramatic downregulation. We also observed downregulation of pathway members downstream of ALK, including in PI3K/AKT and ERK/MAPK pathways as described previously (Miyawaki et al., 2017). Interestingly, we observed differential regulation of PTPN11 upon ALK inhibition. Two C-terminal sites (Y542, Y580) showed marked reduction at 24 hours (Figure 2A); however, the N-terminal PTPN11 site (Y62) was upregulated at 24 hours (Figure 4B). In patient LUAD samples harboring the *ALK* fusion, deep-scale discovery phosphoproteomics (Gillette et al., 2020) also observed outlier phosphorylation of Y542 and Y580 (Figure 2C). These data suggest that ALK inhibition leads to significant downregulation of the C terminal phosphosites on PTPN11, indicating a key role of this phosphatase in ALK-mediated downstream signaling both in cell lines and human tumors. Intriguingly, Gillette et al. also observed a dramatic upregulation of PTPN11 Y62 in *EGFR* mutant tumors (Figure 2D). Consistent with this finding, we observed not only upregulation of Y62 site upon ALK inhibition, but also significant phosphorylation of ERBB3 (Y1289) and EGFR (Y1172) suggesting that activation of alternative receptor tyrosine kinase (RTK) signaling, likely mediated by EGFR and ERBB3 independently or upon their dimerization. This observation, considered in the context of human data, is of significant clinical interest, as resistance to ALK inhibition is often seen in LUAD patients with *ALK* fusion (Dardaei et al., 2018; Rothenstein and Chooback, 2018). The regulation of N- and C-terminal sites on PTPN11 upon ALK inhibition observed using the SigPath targeted phosphopeptide assay, validates and expands on an observation we recently made in the large scale proteogenomic analysis of human LUAD tumors (Gillette et al., 2020).

**Figure 2.**
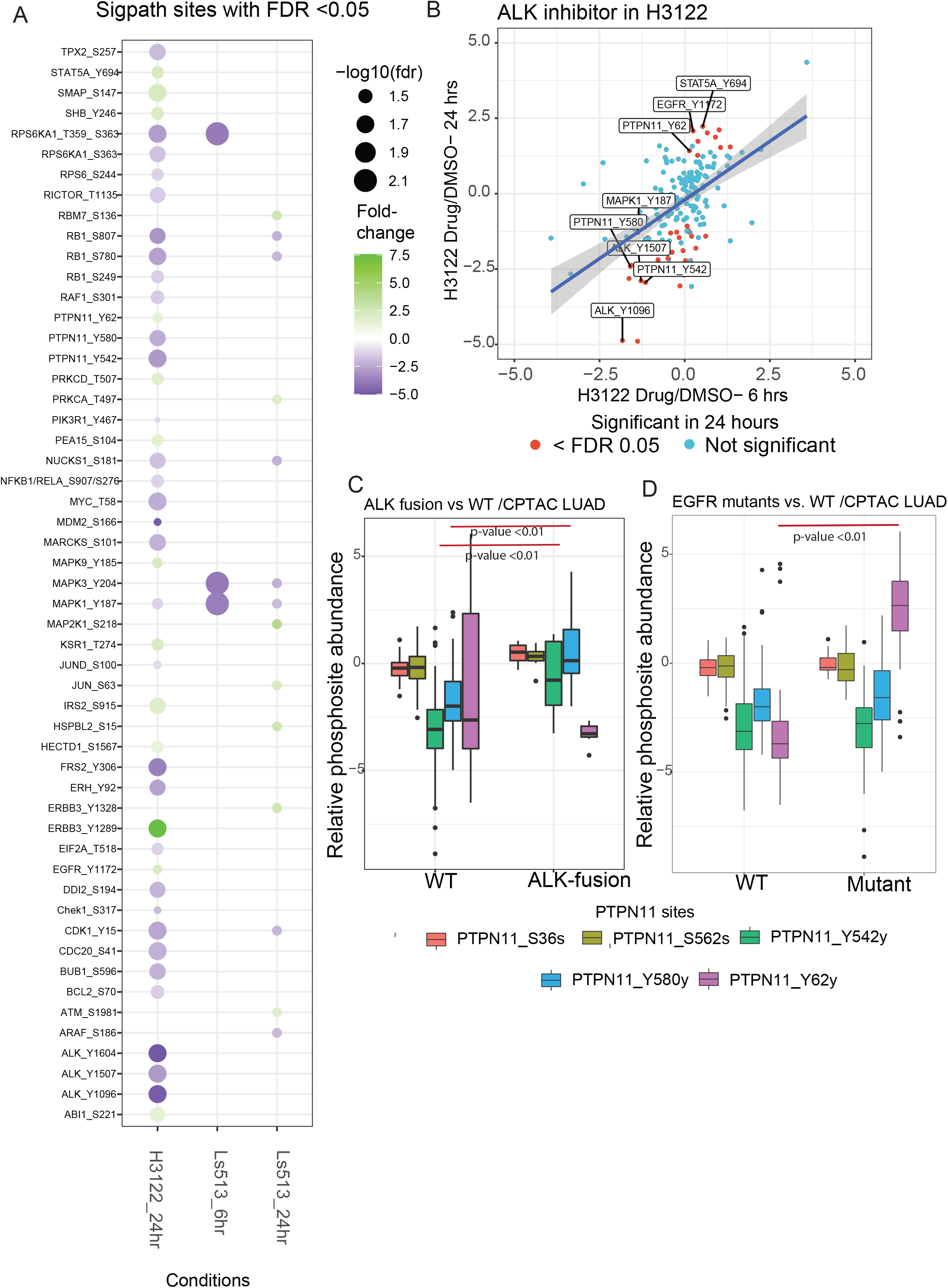
SigPath analyses of cancer cell line perturbation experiments. (A) Summary of all significantly regulated phosphosites observed in H3122 and Ls513 cell lines. The log2 transformed light/heavy peak area ratios for two replicates per time point and treatment were used to compare drug treatment to DMSO. Sigpath phosphopeptides differentially regulated upon treatment in each of the conditions in a moderated two sample T-test (adj. p-value < 0.05) are shown as circles. H3122 6hr experiment didn’t yield any significant regulation and hence not shown on the figure. The color indicates fold change relative to DMSO and the size of the circle indicates log10(FDR). H3122 cell line has no significant regulation at 6 hours and hence is not included in the figure. (B) Scatter plot showing fold-change of Sigpath sites relative to DMSO for H3122 cell line treated with ALK inhibitor. X and Y axis shows 6 hours and 24 hours time points, respectively. The red dots indicate sites with FDR < 0.05. Highlighted are a subset of key differentially regulated phosphosites. Phosphorylation of EGFR, ERBB3 and PTPN11 Y62 are upregulated at 24 hours. (C) Box plot showing relative abundance of detected PTPN11 phosphosites between ALK-fusion containing treatment-naive LUAD patient tumors versus tumors with wild-type ALK. PTPN11 Y546 and Y584 sites show significant differential expression between sample with and without ALK-fusion. These two sites are significantly downregulated in H3122 cell lines upon ALK inhibition. (D) Box plot showing relative abundance of detected PTPN11 phosphosites between EGFR mutated treatment-naive LUAD patient tumors versus tumors with wild-type EGFR. Out of all detected PTPN11 phosphosites, only PTPN11 Y62 shows significant differential expression between EGFR mutant vs. WT samples. The PTPN11 Y62 is upregulated upon ALK inhibition in H3122 cell line, and is likely mediated by the activated EGFR-ERBB3 RTK.

In the Trametinib treated Ls513 colorectal cells we observed strong downregulation at both 6 and 24h of MAPK3 (ERK1) pY204 and MAPK1 (ERK2) pY187, both downstream of MEK consistent with MEK inhibition. Certain drug treatment effects were exclusively observed at either 6 or 24h. For example, downregulation of RPS6K1(pT359 and pS363), which is downstream of ERK, was observed at 6 hours but not 24 hours after Trametinib. Conversely, phosphorylation of RB1 (Y780 and pS807) and CDK1 (Y15) were significant only after 24 hours, suggesting an indirect effect on cell cycle progression. RB1 phosphorylation but not mutation or copy number alteration was shown to drive colon cancer progression (Vasaikar et al., 2019), and our data suggest that Trametinib might downregulate RB1 phosphorylation and subsequently effect cell proliferation.

### Application of the assay to breast cancer xenograft tissue samples

The ability to robustly quantify protein phosphorylation events in tumor samples remains limited. Hence, we sought to test the performance of the SigPath assays in tumor cells undergoing *in vivo* treatment. Previously, the Broad proteomics group and collaborators at Washington University, St. Louis carried out a deep proteome and phosphoproteome study of six patient-derived xenograft (PDX) models of triple negative breast cancer, each carrying unique mutations in the PI3K pathway (Mundt et al., 2018). These PDX models were selected based on their sensitivity to the PI3K inhibitor Buparlasib and exhibited a range of sensitivity to drug treatment with WHIM4 being the most sensitive and WHIM12 the most resistant. After Buparlisib treatment, a clear effect was observed in the phosphoproteome, with downregulation of phosphosites involved in PI3K signaling. However, due to limits on sensitivity and the stochastic nature of data-dependent mass spectrometry-based proteomics especially for modified peptides, both of the canonical AKT phosphorylation sites at threonine 308 and serine 473 were neither quantified nor detected.

To assess the SigPath performance in this context, we applied the full SigPath assay, including pY antibody and IMAC enrichments, to tissue samples from the six PDX models (Figure 3A). 202/298 (68%) phosphopeptides in the SigPath assay were detected at the endogenous level in this study (Table S3). The well-known markers for PI3K inhibition, AKT1S1 pT246, AKT pT308 and AKT pS473 were readily detected and quantified in all models and treatment conditions using the SigPath assay (Figure 3B). These sites were all shown to be significantly downregulated (FDR < 0.05, moderated one-sample T-test) as pharmacodynamics markers of PI3K inhibition following 50hr Buparlisib treatment, with AKT pS473 being significantly downregulated in the 2hr treatment as well. Phosphorylation inhibition of these sites is dependent on the continuous administration of the drug as evident from the upwards trend of the sites following drug washout (Figure 3C). Importantly, the SigPath targeted MS data clearly shows quantitative differences in site modulation between the sensitive and resistant models upon inhibition.

**Figure 3.**
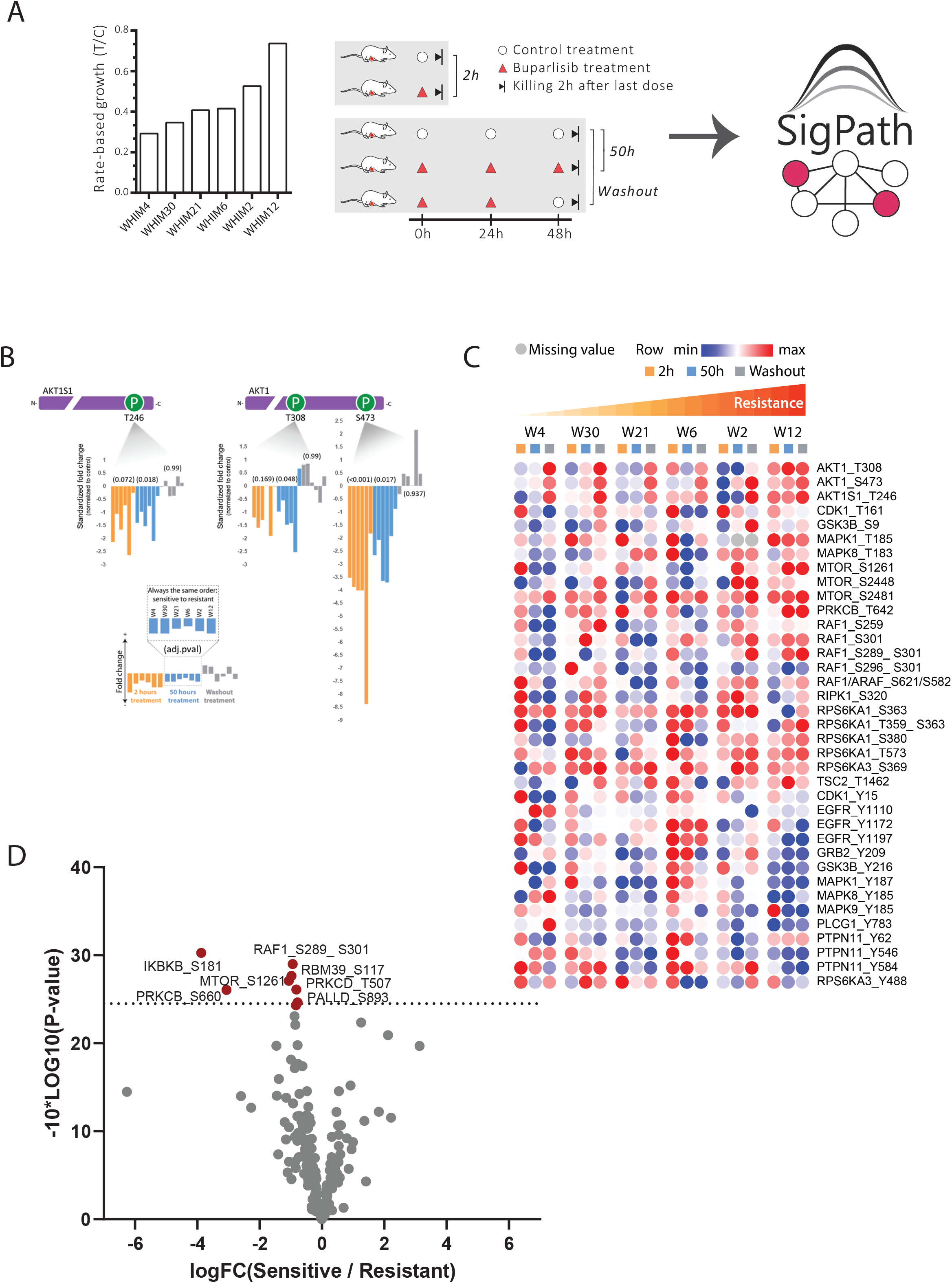
Application of SigPath to understand mechanisms of response and resistance of triple negative breast cancer to therapy. (A) Six patient-derived xenograft models of triple-negative breast cancer were assessed for their resistance to Buparlisib, a PI3K inhibitor, and analyzed for their proteome and phosphoproteome (Mundt et al., 2018). The 6 models ranked after their resistance, from most sensitive to the left (WHIM4), to most resistant to the right (WHIM12). The resistance is calculated as rate-based growth (treatment over control; T/C). Each PDX model was then treated with Buparlisib and tumors were collected at hour 2 or 50 (buparlisib/vehicle administered at hours 0, 24, and 48). A total of 30 samples for the 6 PDX models were analyzed with mass spectrometry-based proteomics and phosphoproteomics. (B) Levels of, AKT1S1 pT246 as well as AKT1 pT308 and pS473 marker sites for Buperlisib activity and PI3K inhibition observed using SigPath. The models are sorted in the order of their resistance to the drug with least resistant on the left. Levels of AKT1 pS473 were reduced in all sensitive models by 80-90%, while this site in the most resistant model (WHIM12) is only reduced about 50%. (C) Heatmap of 37 sites from Hallmark’s PI3K_AKT_mTOR pathway and mTOR detected in Sigpath assay. Ratio of buperlasib treatment to vehicle for each time point is used. WHIMs are listed in the order of their resistance to buperlasib treatment. (C)Volcano plot comparing resistant versus sensitive models in 50hr treatment samples. Sensitive (WHIMs 4, 30, 21, and 6) and resistant (WHIMs 2, and 12) are compared in two-sample moderated T-test. Red dots indicate the 8 peptides significantly regulated with adj. p-value threshold of less than 0.1.

In an effort to investigate pharmacodynamic markers, we looked at the subset of the SigPath data that includes all proteins in Hallmark’s PI3K_AKT_mTOR_signaling pathway and mTOR itself. Of the 48 phosphosites represented in the SigPath assay, 34 were readily quantified across the 6 models and treatment conditions. The ratio of Buparlisib to vehicle treatment for all models listed in the order of their sensitivity to the treatment is shown in Figure 3C. To explore this further and identify response markers, we applied a 2-sample moderated T-test to compare the 2 most resistant PDX models (W12 and W2) with the other 4 models (Figure 3D). Among the highly-upregulated sites in the resistant models are pS289 and pS301 of RAF1, which is regulated by MAPK3 (ERK1) (Balan et al., 2006). The study by Mundt and colleagues. (Mundt et al., 2018) showed that some of the resistance in the most resistant model (W12) was mediated by MAPK3 activation. Our observations using the SigPath assay strengthen the hypothesis that RAF1 is involved in resistance to Buparlasib.

Next we compared SigPath results to the results obtained in a discovery study of the same samples that used chemical labeling with Tandem Mass Tag (TMT) reagents for quantitation of phosphoproteome (Mundt et al., 2018). 146 peptides were quantified in both studies. Comparison of TMT ratios to those in the pooled reference sample with the ratios of endogenous peptide signals to those of the heavy peptides in the SigPath dataset demonstrates the effect of ratio compression in TMT datasets and illustrates how targeted MS overcomes this issue (Figure S3).

### Application of the assay to tumor tissue from medulloblastoma patients

Next, we evaluated human tumor tissue samples with the SigPath assay (Figure S4A). Previously, we carried out deepscale proteomics, phosphoproteomics (including IMAC and pY-enrichment) and acetylproteomics analyses of tissue samples from ca. 40 medulloblastoma patients representing all established subgroups (WNT, SHH, Gr3, and Gr4) (Archer et al., 2018). The study showed that tumors with similar RNA expression vary extensively at the post-transcriptional and post-translational levels. Moreover, proteome profiling revealed additional subgroups within SHH and Gr3 groups, hinting at previously undescribed signaling pathways, as well as providing more prognostic information.

We applied the IMAC subset of SigPath (Table S1A) to the same tissue samples we had previously profiled. The pY subset of SigPath was not applied due to limited sample availability. A total of 138 of the phosphopeptides targeted in the assay (about 60%) were detected across the 38 samples analyzed (Table S4), of which 45 phosphosites were uniquely detected and quantified with the SigPath assay (Figure S4B). The behavior of these 45 sites across all the patient samples is illustrated in Figure 4A. While verification is required, some of the unique sites could constitute novel markers for certain medulloblastoma subtypes. For example, the pS127 phosphosite of YAP1 is upregulated in the sonic hedgehog subtype as compared to both groups 3 and 4 in a one-way Anova test (Figure 4B). The Yap1 protein is amplified and upregulated in hedgehog-associated medulloblastomas (Fernandez et al., 2009) while the quantified Yap1 pS127 site indicates inactivation of the protein in this subtype (Artinian et al., 2015). To investigate whether this can be attributed to a protein-level difference, we extracted protein ratios from the proteomic discovery study for all patient samples and performed correlation analysis with the SigPath assay ratios. High correlation (R2 = 0.69) indicates that in our SigPath assay YAP1 pS127 is acting as a proxy for the protein level difference in this case. In contrast, the pS89 site of TAZ upregulated in sonic hedgehog versus group 4 is not due to protein level difference, as correlation of this phosphosite to the protein is very low (R2 =0.003) (Figure 4B). High expression of the transcriptional coactivator TAZ has been shown to be associated with a worse prognosis and affects cell proliferation in patients with medulloblastoma, regardless of subtype (Wang et al., 2019); however, phosphorylation of the S89 site will lead to an inhibition of carcinogenesis and cell growth (Cordenonsi et al., 2011; Zhang et al., 2015). These new phospho-level findings may be of significant interest to the medulloblastoma community.

**Figure 4.**
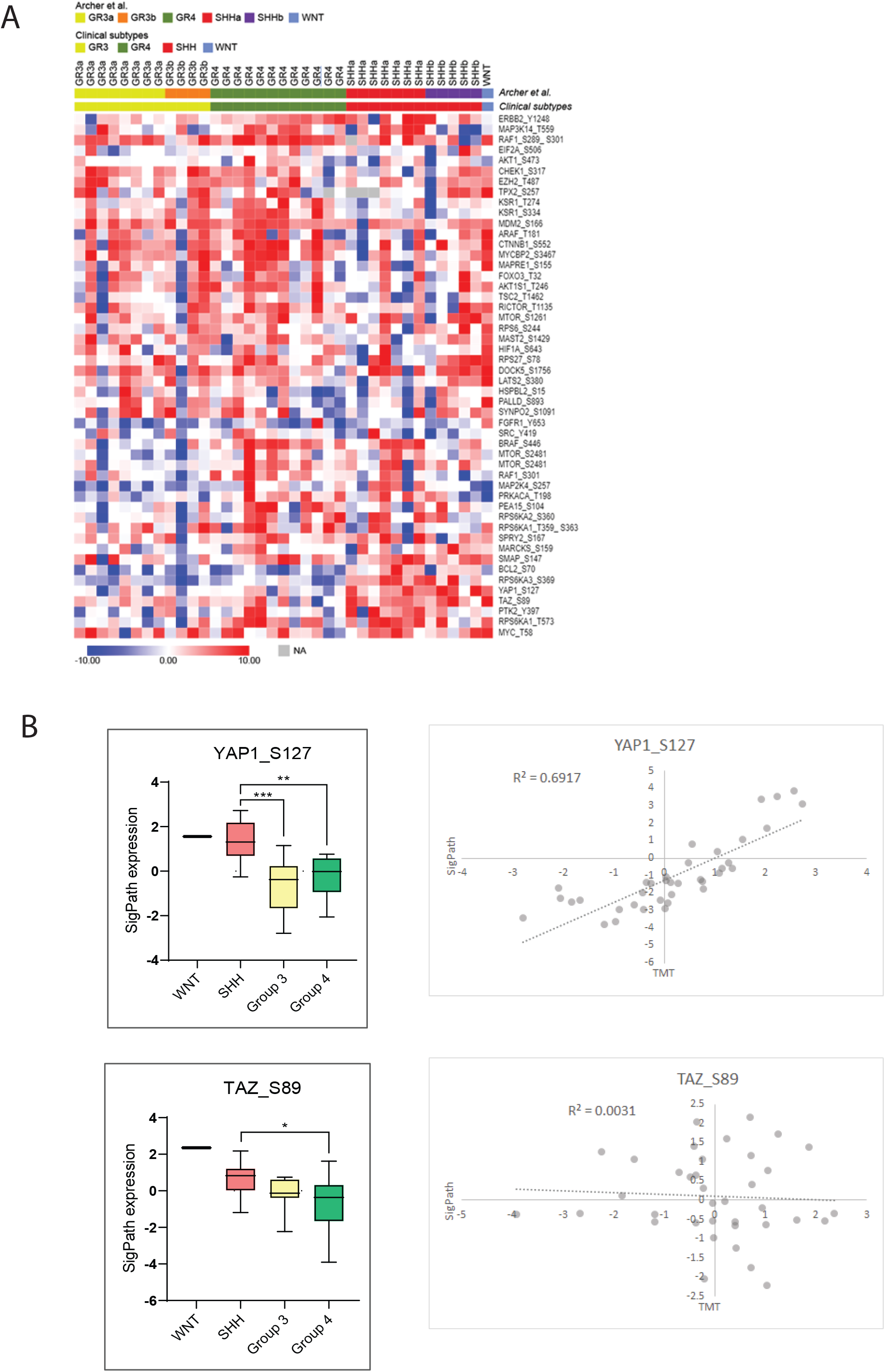
Application of the IMAC subset of SigPath to cancer tissue samples from 37 medulloblastoma patients. These patients represent all the known clinical subtypes; WNT (n = 1), SHH (n = 12), group 3 (GR3; n = 12) and group 4 (GR4; n = 12). (A) A heat map of 37 patients with medulloblastoma show how 45 phosphosites uniquely detected in SigPath assay. Samples are clustered by their original clinical subtypes as well as by new classification in Archer et al. (Archer et al., 2018) where discovery analyses split subgroup 3 into 3b and 3a, and subgroup SHH into SHHa and SHHb. The heat map was generated using Morpheus online tool. (B) Box plots comparing all the data for YAP1 pS127 and TAZ pS89 for the different groups of medulloblastoma (SHH, GR3, GR4 and WNT). One-way ANOVA with an *ad hoc* Tukey’s test (with adj. p-values for multiple comparisons) was applied for the comparison. Scatter plots comparing TMT ratios for YAP1 and TAZ proteins to SigPath ratios for YAP1 pS127 and TAZ pS89, respectively. Pierson correlation coefficient is shown on the plots.

We also looked at the overlap of the 138 phosphosites detected in the SigPath assay with those from the discovery data (Archer et al., 2018) and found that 61/138 peptides were detected in all samples in the discovery dataset, while another 27 were detected in at least one sample of the discovery dataset. Correlation analysis of discovery and SigPath results for the 88 sites detected in both demonstrated a high level of agreement between phosphosite abundances as measured by both platforms (Figure S4C).

## Discussion

Targeted MS assays have largely been developed for purposes of biomarker verification, but are increasingly being used to validate findings in biological and preclinical studies (as reviewed in (Parker and Borchers, 2014; Rifai et al., 2006). The highly multiplexed, targeted quantitative assay we developed and applied here was designed to measure phosphosites in nodes of biological pathways known to be modulated via phosphorylation including the RAS, MAPK, PI3K, PKC, SRC and JAK signaling pathways, transcription factor circuit nodes including FOXO, STAT, NFKB, TGFB and Wnt pathways, as well as protein phosphosites useful as readouts of DNA damage, cell cycle arrest, apoptosis, spindle checkpoint activation, hypoxia, autophagy, cell stress and epithelial-to-mesenchymal transition. Through a range of applications in cell lines, preclinical models and clinical samples, we demonstrated the utility of the assay to not just verify prior findings, but to detect and quantify a large number of differentially regulated phosphosites newly associated with drug perturbation and disease subgroups. For example in Alk fusion cell line with the treatment of Ceritinib we observed differential phosphorylation of N- and C-terminal phosphosites from PTPN11 phosphatase consistent with observations in LUAD tissue samples (Gillette et al., 2020). These results suggest that SigPath, and targeted MS assays in general, should be more routinely used in the development of optimization of therapeutics. Our results also highlight the potential of SigPath to monitor phosphoproteomic signaling events and to nominate mechanistic hypotheses regarding oncogenesis, response and resistance to therapy in disease models and human tumors.

The ca. 300-plex SigPath assay was designed to flexibly allow measurement of phosphosites having phosphotyrosine alone, chiefly phosphoserine- and phosphothreonine-sites, or a mix of all three. Any proteomics lab skilled in phosphopeptide enrichment can implement the assay, and phosphopeptide sample enrichment is readily automated (Abelin et al., 2016). The SigPath assay is readily extendable to measure other phosphosites by synthesis of the new phosphopeptides in heavy-labeled form and repeating the QC processes described with a focus on the new sites. Three hundred targets does not represent the maximal level of multiplexing that can be achieved. Using modern MS instrumentation and techniques such as internal standard-triggered parallel reaction monitoring (Gallien et al., 2015), assay panel sizes can be increased to 500 targets or more. In contrast, assays employing anti-peptide antibodies (Keshishian et al., 2015; Kuhn et al., 2009; Kuhn et al., 2012; Sperling et al., 2019; Whiteaker et al., 2014; Whiteaker et al., 2011) take a long time to develop and qualify, are very costly even for small numbers of targets, have yet to be multiplexed to the level demonstrated here and are currently available in only a few expert labs, limiting their utility for the biology community. A limitation of SigPath as configured is that it profiles phosphorylation only, and therefore cannot assess whether the observed regulation of phosphorylation was due to an altered protein expression or a post-translational effect. It could be possible, depending on overall protein abundance, to also measure global protein expression and hence derive phospho-stoichiometry by adding heavy non-phosphorylated peptide standards and measuring them in the post-IMAC flowthrough in this workflow.

While Western-competent antibodies exist for many of the targets in SigPath, they are generally deployed in a highly selective and individual manner and are non-quantitative, resulting in large swaths of biology remaining opaque to the investigator. Multiplexed Western and ELISA panels are commercially available, but typically measure only 10-20 analytes per well in order to avoid cross-reactivity and maintain sensitivity (e.g., [Quanterix](https://www.quanterix.com/), [MSD](https://www.mesoscale.com/en/site_search?keyword=phospho&manual-search=true), [Myriad](https://myriadrbm.com/search/?addsearch=phospho), [Abcam](https://www.abcam.com/products?keywords=phospho), [AbcamReview](https://www.abcam.com/kits/multiplex-immunoassay-techniques-review-of-methods-western-blot-elisa-microarray-and-luminex#single)). Large protein assay panels employing two different antibodies for each protein target with detection based on proximity extension are also now commercially available from Olink ([Olink](https://www.olink.com/)). However, these assay panels primarily measure proteins, not phosphosites.

The depth of detection in any targeted MS approach is governed by the abundance of the target peptide in the sample being analyzed, the efficiency of the enrichment process and the amount of input peptide that was enriched. When IMAC phosphopeptide enrichment alone is used, the efficiency is ca. 95%, that is, less than 5% the peptides observed after enrichment do not contain a phosphorylated amino acid. This is important, as non-phosphorylated peptides are generally present at much higher abundance than the phosphopeptides and so can interfere with detection of phosphopeptides. In the studies presented here, 500 to 1000 μg of peptides were used for the IMAC portion of the assay; however as little as 50 μg can be used, with consequent loss of detection of a subset of the least abundant phosphopeptides (data not shown). Due to the lower abundance of phosphotyrosine-containing peptides, antibody-based phosphotyrosine peptide enrichment typically requires a larger sample input, often in the range of 1-5 mg, to get high coverage of the sites in the SigPath assay.

Once targeted MS-based assays achieve the large numbers presented here in SigPath, they can also be viewed as a different, complementary way to do discovery, with a narrower spectrum but more consistent measurements and vastly greater throughput. For example, the bona fide PI3K inhibition marker AKT S473 was missed completely in the PDX discovery data set, while the SigPath assay consistently quantified this site. Sample preparation for SigPath is much simpler than methods commonly used for deep-scale discovery, requiring only digestion, phosphopeptide capture and analysis of the captured peptides together with spiked heavy peptide standards. In addition, the on-instrument time is also 6-12x faster than discovery methods that require fractionation of the peptides off-line prior to LC-MS/MS to gain depth of detection. Quantification of phosphopeptides using SigPath is far more precise than label-free discovery experiments or those using chemical labels like TMT, as targeted MS methods do not suffer from the ratio compression challenges of the latter. While the breadth of coverage in SigPath is much smaller than discovery phosphoproteomics analyses (>300 phosphopeptides compared to >30,000 phosphopeptides per sample), the quantitative precision and repeatability for targets measured by the assay confer their own advantages, as shown here in the replicates of the ALK inhibitor study and in other phosphopeptide assays we have uploaded to the CPTAC assay portal. Since SigPath detects and measures the spiked heavy peptide forms of all of the phosphopeptides targeted, determining the presence and level of the endogenous form by ratioing the intensities of the endogenous and spiked peptide-specific fragment ions, it, like other targeted MS assays, is far less susceptible to false negatives: if the target was not detected, it was below detection limits. Therefore, we view SigPath as an impactful resource for the cancer research community, suitable both for discovery, as a verification assay for targets of biological import and for preclinical studies in human cancers and other diseases.

## Acknowledgments

This work was supported by a grant from Novartis and partially by NIH/NCI grants from the National Cancer Institute (NCI) Clinical Proteomic Tumor Analysis Consortium grants NIH/NCI U24-CA210986 and NIH/NCI U01 CA214125 (to SAC) and NIH/NCI U24CA210979 (to DRM). We thank Jacob Jaffe for help with searching phosphopeptides against existing datasets.

## Author Contributions

Conceptualization, HK, ERM, DP, KW, WRS, SAC; Methodology, HK, ERM, DP, JJ-V, KW, SAC; Software, KK, DRM, KC; Validation, HK, RM, LW, MB, MM, DRM; Formal Analysis, HK, KK, FM, SS, RM, LW, HS, EK, DRM, KC; Data Generation, HK, RM, LW, HS, FM, MB, MM; Resources, ERM, DP, JJ-V, SS, MG, WRS, SAC; Data Curation, HK, ERM, DP, FM, KK, JG, SS, DRM, MG, KC, TR; Writing Original Draft, HK, RM, FM, KK, SAC; Review and Editing of Manuscript, HK, ERM, FM, KK, EK, SS, MG, TR, KW, WRS, SAC; Supervision, HK, ERM, KK, JJ-V, KW, WRS, SAC; Funding Acquisition, WRS, LAG, SAC.

## Main and Supplemental Figure Legends

### Supplemental Figure Legends

**Figure S1:**
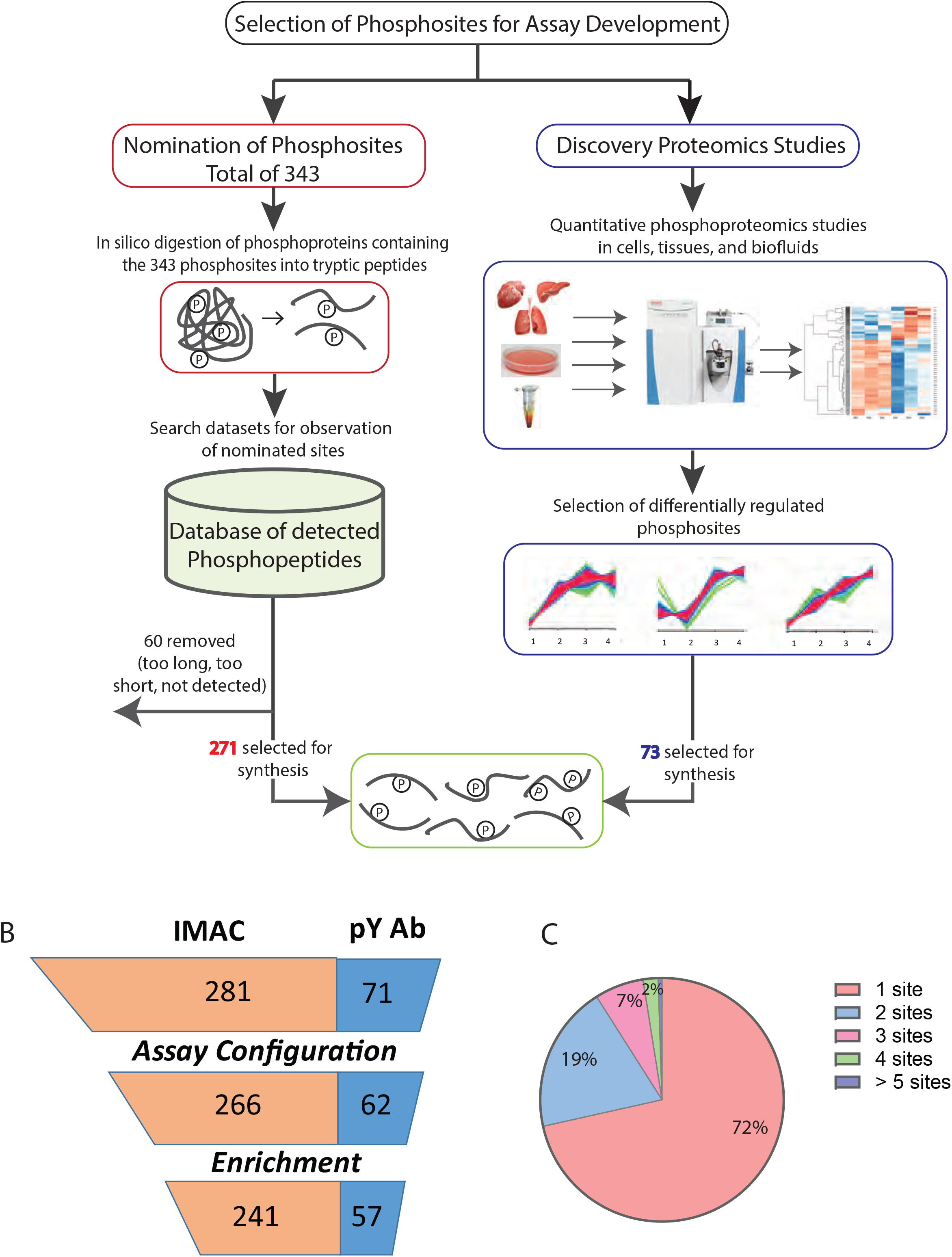
Development of SigPath assay. (A) Process for selecting phosphosites and phosphopeptides for SigPath assay development. Majority of phosphosites were nominated by experts, then converted into tryptic peptides, searched against existing datasets at the Broad for detection of them in MS data. One fourth of the phosphosites were included based on them being differentially regulated in quantitative phsophoproteomics studies (Mertins et al., 2014). Once finalized [C13, N15] labeled version of the peptides were synthesized for the assay. (B) Assay Configuration and testing statistics of SigPath. Twenty four out of 352 peptides failed the assay configuration due to their LC or MS characteristics, while another 30 failed during the pY or IMAC enrichment step. Final SigPath assay targets 298 phosphopeptides with 284 phosphosites. (C) Pie graph showing range of phosphosites per protein in the assay panel. 72% of the proteins are represented by only one phosphopeptide, 15% by two phosphopeptides. The remaining varies from 3 to 9 phosphopeptides.

**Figure S2:**
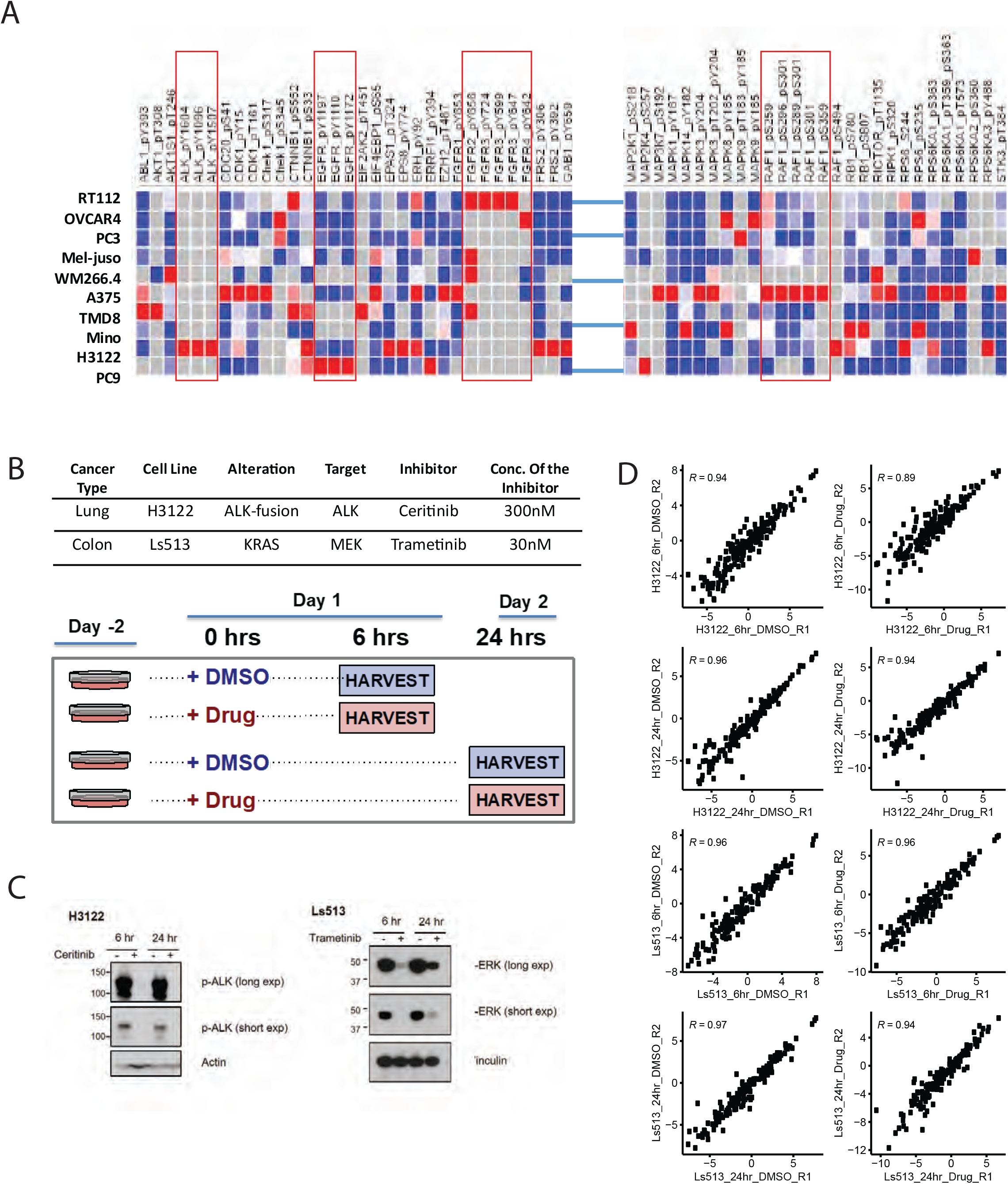
Application of the assay to cell line samples at the baseline (A) Heatmap showing subset of the results across the 10 cell lines. Endogenous light to SIL heavy peptide ratio is used for the plot after row normalization (B) Experimental design and details for drug treatment studies in H3122 and Ls513 cell lines. Table contains details about the cell lines as well as the inhibitor and concentration of it used. Cells were treated either with the inhibitor or DMSO for 6hours and 24hours. Two process replicates were collected for each sample (C) Inhibition of pALK and pERK signaling in established human cell lines. Immunoblot analyses of cultured H3122 lung adenocarcinoma cells (on the left) treated with ALK inhibitor Ceritinib (+) or DMSO (−) for 6 and 24 hours. Antibodies recognizing the phosphorylated 1507-Tyrosine site of the ALK protein and the Actin protein (loading control) were used. Immunoblot analyses of cultured Ls513 colorectal carcinoma cells (on the right) treated with KRAS inhibitor Trametinib (+) or DMSO (−) for 6 and 24 hours. Antibodies recognizing the phosphorylated Thr 185/Tyr 18) sites of the ERK1/ERK2 proteins and the Vinculin protein (loading control) were used. (D) Scatter plot of two process replicates of Log2 light/heavy PAR of each sample group. Shown on each plot is the Pearson correlation coefficient.

**Figure S3:**
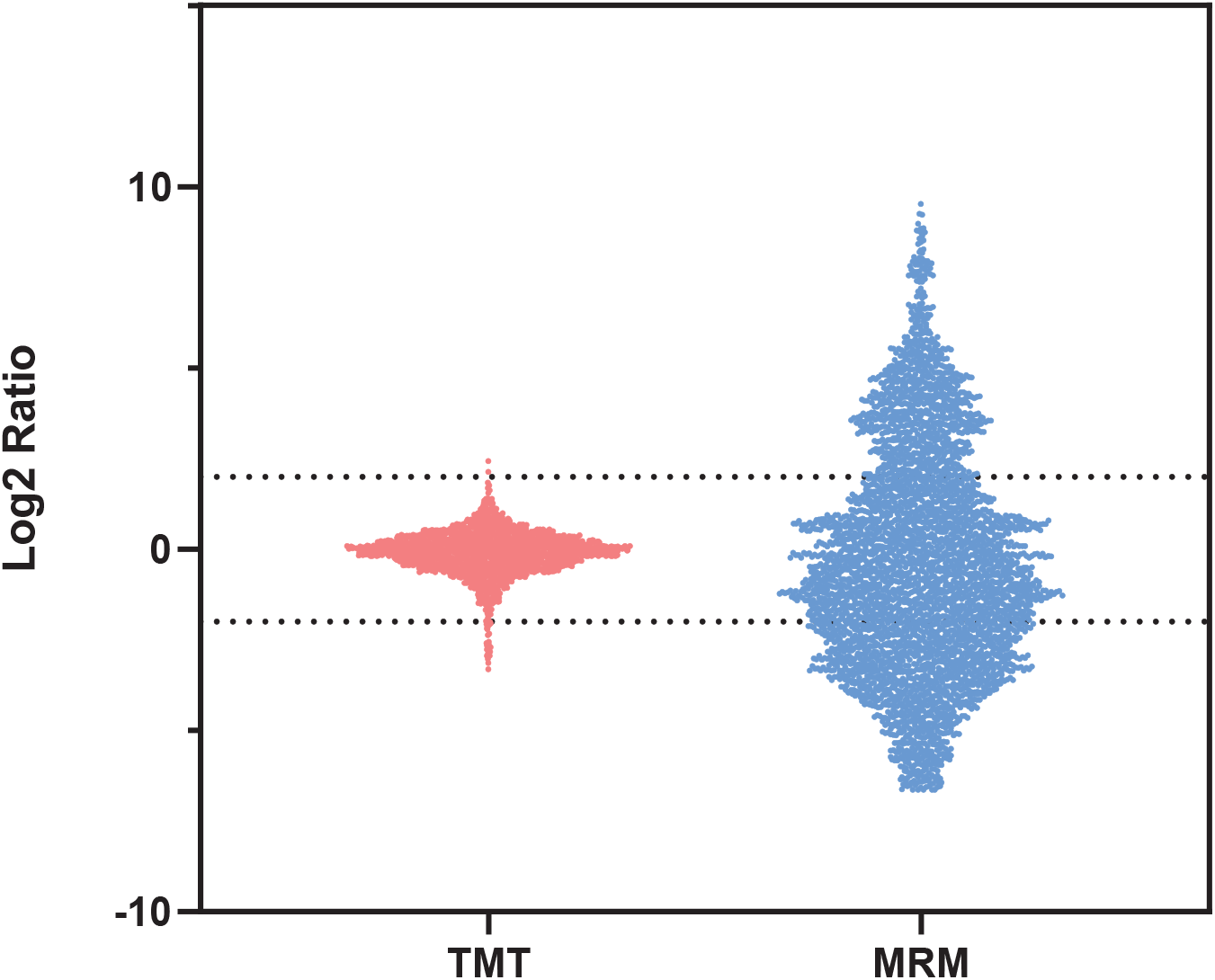
Bar and whisker plot showing the range of ratios obtained for the overlapped 146 peptides in discovery study using TMT and SigPath. For discovery study Log2 ratio of all the peptides to the pooled reference was used for the plot. For SigPath Log2 ratio of light to heavy peptide was used for the plot. Any ratio less than 0.01

**Figure S4:**
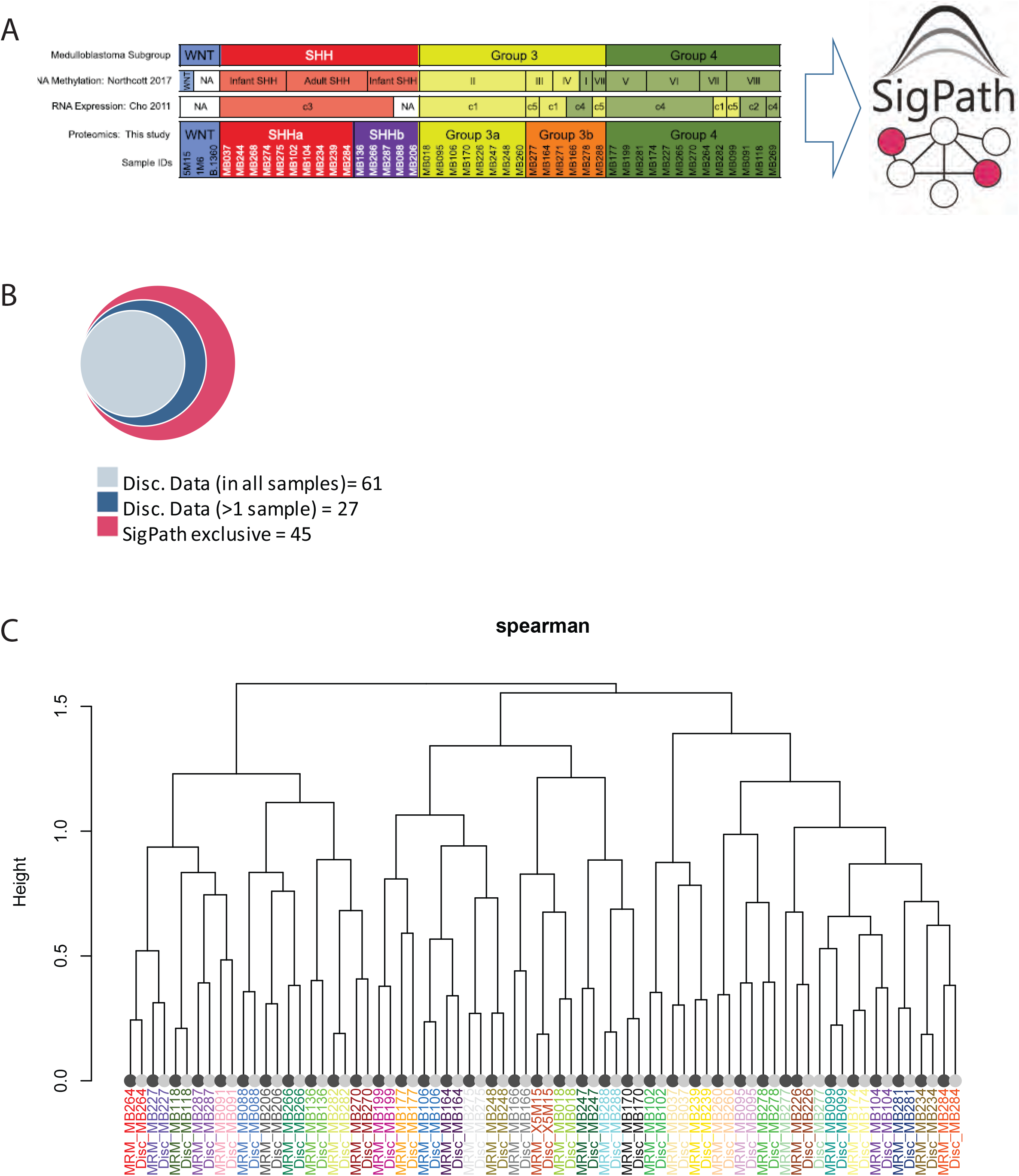
Application of the IMAC subset of the SigPath assay to human tissue samples from medulloblastoma patients (A) List of samples used for this experiment and their classification to SHH, GR3 and GR4 as well as subgroups within GR3 and GR4 ((Archer et al., 2018). Only IMAC subset of the assay has been applied to these samples (B) Venn diagram showing the overlap of the 134 sites detected in SigPath assay with the discovery data. 61 peptides were detected in the discovery dataset in all samples (gray) while another 27 detected in at least 1 sample (blue) of the discovery dataset. 45 sites (30%) were unique to SigPath assay (red). (C) Dendrogram illustrating the clustering of MRM (dark leaves) and discovery data (light leaves) of 88 phosphosites detected in both assays. Colors of sample identifiers are coordinated by patient. All but one sample pair (MRM/discovery) cluster adjacent to each other. Dendrogram was derived from complete-linkage hierarchical clustering using 1-Spearman correlation as the distance metric.

## Supplemental Table Legends

**Table S1A: List of synthetic stable isotope labeled (SIL) peptides for SigPath assay**. List includes all the peptides that were synthesized sorted by the gne symbol. Red amino acids in the Phosphopepide Sequence column indicate [C13, N15] labeled amino acid. *Failed peptides column indicate all the peptides that failed either during assay configuration due to LC and MS characteristics or pY or IMAC enrichment step. Blank cells indicate working peptides.

**Table S1B: Pathway coverage with the working SigPath assay.** Pathway coverage is assessed using all the canonical databases in MSigDB 3.0. Table includes number of genes, phosphosites and phosphopeptides in SigPath panel for each of the pathways

**Table S1C: List of phosphosites nominated but not considered for SigPath assay.** List includes the sites, tryptic peptides containing the sites and reason for not including in the panel.

**Table S2A: List of 10 cell lines used for assessment of the SigPath assay.** Table includes details about the cell lines t and number of peptides quantified for each of them.

**Table S2B: Data table of perturbagen experiment results in H3122 and Ls513 cell lines.** Table includes all the peptides detected in both experiments with results of moderated two-sample T-test. Peptides are identified by the gene symbol and phosphosite.

**Table S3: Data table of all PDX samples.** Log2 Light/Heavy peak area ratios (PAR) of all the phosphosites quantified in all the time points of PDX models.

**Table S4: Data table of all medulloblastoma samples.** Log2 Ligth/Heavy PAR of all the phoshphosites quantified in all medulloblastoma samples.

## Methods and Protocols

### Assay development

#### Peptide selection by nomination and selection from phosphoprofiling studies

For the list of nominated phosphosites the first step was to convert them into tryptic peptides containing the sites .An *in-silico* tryptic digest of the Uniprot protein database was generated to identify fully cleaved tryptic peptides containing the nominated phosphosites. Peptides containing more than 40 amino acids in the list were dropped at this step of the selection process. Too short fully cleaved tryptic peptides (< 6 amino acids) were still considered at this step in case missed cleaved versions with reasonable length were observed in the existing datasets (Figure S 1A).

Next step was to ascertain whether they had been detected by mass spectrometry. For this purpose, we used the very large collection of high quality phosphopeptide data generated by the proteomics group at the Broad Institute over the past 15 years to identify whether and in which form(s) the tryptic phosphopeptide had been most frequently observed (e.g., full tryptic, incomplete cleavage, etc.). For the search we utilized in-house developed R and Perl scripts specifically tailored towards database search results created by Spectrum Mill Software (Agilent Technologies, Santa Clara, CA) or MaxQuant (Cox and Mann, 2008). Only validated peptides at 1% false discovery rate (FDR) were considered in subsequent analysis. Miss-cleaved versions of the query peptide were allowed. Peptides identified by this approach were manually analyzed for each peptide to select a version for synthesis. In the cases where more than one version of the peptide was observed (fully cleaved or missed cleaved, singly or multiply phosphorylated) priority was given to the version with the highest frequency of observation in the datasets. In selecting singly versus multiply phosphorylated version of a peptide, priority was given to singly phosphorylated version unless doubly phosphorylated version was much more prevalent in existing datasets. Moreover, for MAPK1, MAPK14, MAPK3, MAPK8, MAPK9, RAF1, and RPS6KA1 in addition to a singly phosphorylated version also included doubly phosphorylated forms of the peptides (see Table S 1A). There are no triply phosphorylated peptides in the assay panel.

Seventy percent of sites nominated (234 out of 343) were previously observed as fully tryptic phosphopeptides or in a mis-cleaved form in our experimental datasets. Table S 1A lists all of the previously observed nominated phosphosites, the phosphoprotein of origin, the dominant tryptic peptide form containing each site and the enrichment methodology required for detection of the site (i.e., either immobilized metal affinity chromatography (IMAC) or anti-phosphotyrosine (pY) antibody). Sixty-two phosphopeptides containing 66 of the 343 nominated phosphosites were not observed in our datasets (Tables S 1A and S 1C). Despite the lack of prior observation in discovery data, we included 37 of these phosphopeptides due to their importance in cancer biology and the potential for the targeted MS method to have greater sensitivity for their detection than the discovery methods used. The remaining 29 sites were excluded from consideration (Table S 1C).

In addition to lack of detection in experimental data, there were a number of other reasons to not advance to assay configuration for some of the nominated phosphosites, including target peptides being too long or too short or having failed synthesis. Nineteen of the nominated sites were located in tryptic peptides that were deemed to be too long (>40 amino acids) because of anticipated issues with synthesis, chromatography and/or assay development and were eliminated (Table S 1C). In addition, a number of nominated sites were located in short tryptic peptides of 6 amino acids or less. To improve likelihood of detection, specificity and chromatographic retention on C18 reverse phase matrix, we instead searched the data for longer, mis-cleaved forms of these peptides. Seventeen of the total 25 short peptides were found in longer, mis-cleaved peptides (>6 but <40 amino acids) and were included for assay development. For example, the nominated site S380 in Serine/threonine-protein kinase LATS2 is present in tryptic peptide form D(pS)LQK. The longer, mis-cleaved version of this peptide RD(pS)LQKPGLEAPPR was found in the discovery data and selected for assay configuration (Table S1A). Mis-cleaved forms for eight additional short phosphopeptides were not detected in the data and so these were removed for further consideration (Table S 1C).

Seventy-three phosphosites in the panel were derived from quantitative proteomics discovery studies in our laboratory and were among the most significantly regulated sites in those studies. 17 phosphopeptides were selected from a discovery phosphoprofiling study investigating impact of ischemia in ovarian cancer and breast cancer xenograft tissues (Mertins et al., 2014).

The remaining experimentally derived sites were included based on the analysis of several discovery experiments done in cancer cell lines treated with specific inhibitors.

During the selection process, the uniqueness of each peptide in the human proteome was also taken into consideration with priority given to peptides unique to one protein. Seven of the peptides in the final assay panel are shared with more than one protein (Table S1). Among these are: pS909 and pT1079 of LATS1; pS872 and pT1041 of LATS2; pT35 of MOB1A and MOB1B; pS907 of NFKB1 and pS276 of RELA; pT198 of PRKACA, PRKACB, and PRKACG; pS621 of RAF1 and pS582 of ARAF: and pY706 of NTRK2 and pY709 of NTRK3.

### Peptide synthesis

The final list of 352 tryptic phosphopeptides representing 344 phosphosites (17 peptides having more than one site) corresponding to 234 phosphoproteins were synthesized containing single stable isotopically labeled (SIL) amino acid. Most peptides contained [13C, 15N] lysine or arginine at the C-terminus. Nine peptides representing C-terminal of the protein were synthesized with either heavy N-terminal lysine or arginine, or heavy internal leucine and proline.

### Peptide organization

Synthetic heavy labeled internal standard peptides were organized in mixtures first by their enrichment methodology (immobilized metal affinity chromatography (IMAC) and/or phosphotyrosine (pY) antibody). 231 peptides containing phosphoserine (pS) or phosphothreonine (pT) were organized in five IMAC mixtures alphabetically by gene name containing 43 to 50 phosphopeptides each. 71 pY containing peptides were organized in 2 pY mixes alphabetically by gene name, each containing 31 or 40 peptides. Finally the remaining 50 peptides (predominantly pY) with previous detection information after IMAC as well as pY antibody enrichments were organized in a separate mixture (IMACpY) and used in combination with either IMAC or pY mixtures. Peptides were at 2pm/ul equimolar concentration in all of the mixtures

### Assay configuration

Multiple reaction monitoring (MRM) assay configuration using heavy labeled synthetic peptides was done on TSQ Quantiva triple quadrupole mass spectrometer (Thermo Fisher) coupled with Easy-nLC 1200 ultra-high pressure liquid chromatography (UPLC) system (Thermo Fisher) in several batches of 50 to 100 peptides each. Skyline Targeted Mass Spec Environment was used throughout assay configuration and all data analysis. First, spectral libraries for the peptides were generated on a Q Exactive mass spectrometer. Spectral libraries were uploaded to Skyline and 5-10 most intense fragment ions (transitions) for each peptide were selected for MRM assay configuration. Transitions containing phosphosite or helping with the assignment of the phosphosite were included in the transition list for assay configuration. Next, collision energies (CE) were optimized for all the transitions and peptides by liquid chromatography – multiple reaction monitoring mass spectrometry (LC-MRM/MS) on TSQ Quantiva using Skyline’s CE optimization module. For every transition starting with the instrument specific calculated CE tested 10 additional CEs (5 below and 5 above the calculated CE) in increments of 2. The list of transitions with varying CE values was exported from Skyline and used for building MRM method in Xcalibur software. Equimolar mixture of peptides at 50fm/ul was analyzed by LC-MRM/MS on Quantiva using this method. Resulting data was analyzed on Skyline which then selected the CE that resulted in the highest peak area for each transition. In the final step of CE optimization MRM data was acquired with optimized CE values for every transition. Using this dataset in Skyline, manually selected the best 3 to 6 transitions for every peptide giving highest priority to fragment ions of y-series with mass to charge (m/z) above the precursor, and ions containing phosphosite or helping with the site localization. For the peptides where the options were more limited also included ions of y-series with m/z below the precursor and b-series.

After the CE optimization compiled 2 LC- MRM/MS methods; one for peptides enriched by IMAC strategy (231 IMAC and 50 IMACpY mixtures) and the second for peptides enriched by pY antibody strategy (71 pY and 50 IMACpY mixtures).

Liquid chromatography was performed on a 75um ID picofrit columns packed in-house to a length of 28-30cm with Reprosil C18-AQ 1.9um beads (Dr Maisch GmbH) with solvent A of 0.1% formic acid (FA) / 3% acetonitrile (ACN) and solvent B of 0.1% FA / 90% ACN at 200nL/min flow rate. Below are the details of the IMAC and pY LC-MRM/MS methods:

#### IMAC LC-MRM/MS method

method duration – 160min, gradient – 2 to 6% solvent B in 1min, 6 to 30% B in 124min, 30 to 60% B in 9min, 60 to 90% B in 1min, followed by a hold at 90% B for 5min, and subsequent hold at 50% B for 19min. MS parameters include 3sec cycle time, Q1 and Q3 resolution of 0.4 and 0.7, respectively, retention time (RT) scheduling window of 10min.

#### pY LC-MRM/MS method

method duration – 120min, gradient – 2 to 6% solvent B in 1min, 6 to 30% B in 84min, 30 to 60% B in 9min, 60 to 90% B in 1min, followed by a hold at 90% B for 5min, and subsequent hold at 50% B for 19min. MS parameters include 1.5 sec cycle time, Q1 and Q3 resolution of 0.4 and 0.7, respectively, RT scheduling window of 10min.

### Cell lysis and digestion

Lysis buffer (8 M urea, 75 mM NaCl, 50 mM Tris pH 8.0, 1 mM EDTA, 2 ug/mL Aprotinin, 10 ug/mL Leupeptin, 1 mM PMSF, 10 mM NaF, 1:100 phosphatase inhibitor cocktail 2, and 1:100 phosphatase inhibitor cocktail 3) was added on top of the cell pellets, vortexed lightly, and incubated on ice for 15 minutes. The samples were then vortexed for 10 seconds on the highest setting, and allowed to incubate on ice again for 15 minutes. Following lysis, the samples were spun down at 20,000 rcf for 10 minutes at 4°C to pellet insoluble cell debris. Each cell pellet lysate was then transferred to a new 2 mL Eppendorf tube and quantified by the Pierce BCA Protein Assay Kit (ThermoFisher, 23225). After quantification, the concentrations were all normalized to the lowest sample with lysis buffer, then reduced with 5 mM dithiothreitol (DTT, Pierce, A39255), mixing at 800 rpm for 45 minutes at room temperature. Once reduced, the samples were alkylated using 10 mM iodoacetamide (IAA, Sigma-Aldrich, 144489) for 45 minutes in the dark at room temperature. All lysates were diluted 1:4 with 50 mM Tris HCl pH 8.0 and digested with LysC (Wako, 12505061) at an enzyme to substrate ratio of 1:50 for 2 hours at 30°C and shaking at 800 rpm. Following LysC, trypsin (Promega, V5111) was added at an enzyme to substrate ratio of 1:50 overnight at 37°C and shaking at 800 rpm. To quench the digestion, 10% formic acid was added to a final concentration of 1% and pH 3.

### Peptide cleanup by cartridge desalt

200 mg (3 cc) Sep-Pak C18 Vac Cartridges (Waters, WAT054945) were conditioned with 3 mL of acetonitrile followed by 3 mL of 0.1% FA / 50% ACN. The cartridges were then equilibrated with four 3 mL injections of 0.1% trifluoroacetic acid (TFA). The samples were then loaded onto the cartridges and flowed through. After loading the sample, the cartridges were washed three times with 3 mL 0.1% trifluoroacetic acid and one time with 3 mL 1% FA. The peptide samples were eluted off the cartridges with two injections of 1.5 mL 0.1% FA / 50% ACN into 15 mL falcon tubes. All digests were frozen, dried down, and reconstituted in 0.1% FA / 3% ACN and quantified by the Pierce BCA Protein Assay Kit. Each sample was then aliquotted into 5 mg for pY antibody enrichment, frozen, and dried down by speedvac. If pY antibody enrichment was skipped then 500ug aliquots were made for each IMAC enrichment step (see below).

### Phosphotyrosine enrichment

Peptide aliquots for pY enrichment were reconstituted with 1.5 mL of IAP buffer (50 mM MOPS/NaOH pH 7.2, 10 mM Na2HPO4, 50 mM NaCl) and kept on ice throughout the experiment. Once reconstituted, 30 fmol of the pY and IMACpY heavy peptide mixtures were spiked into each sample, then vortexed, and spun down at 5000 rcf for 5 minutes. The pY 1000 Immunoaffinity beads (Cell Signaling Technology, 8803S) were suspended in 1.5 mL IAP buffer and transferred to a 1.5 mL eppendorf tube. The beads were spun down at 1500 rcf for 1 minute and the supernatant was removed. This washing process was repeated three times through. After removing the IAP buffer supernatant from the last wash, the reconstituted peptide samples were transferred onto the Immunoaffinity beads and mixed end-over-end at 4°C for 1 hour. After the hour incubation, the beads were spun down at 1500 rcf for 1 minute and the supernatant was collected as the pY flow through for IMAC enrichment. The pY 1000 beads were washed four times with 1.5 mL cold phosphate buffered saline (PBS, ThermoFisher, 10010023). After washing, the beads were resuspended with 50 uL of 0.15% TFA and incubated at room temperature for 5 minutes. The beads were spun down and the supernatant was transferred onto a prewashed and pre-conditioned stagetip (see below). The TFA incubation was repeated one more time for a total of 100 uL of supernatant transferred onto the stagetip.

### Phosphotyrosine enrichment stagetip desalt

Stagetips, prepared with two Empore C18 (3M, 2315) punches, were conditioned with 100 uL of methanol and spun down at 3100 rcf for 1 minute. The tips were further conditioned with 100 uL of 0.1% FA / 50% ACN, then equilibrated with 2 injections of 100 uL 0.1% FA. After equilibration, the two 50 uL pY enriched incubations were added to stagetips then spun down. The samples were desalted by spinning down two injections of 100 uL 0.1% FA. The bound peptides were eluted off the stagetips using 50 uL of 0.1% FA / 50% ACN. The eluates were transferred to autosampler vials, frozen and dried down, then reconstituted in 5 uL of 0.1% FA / 3% ACN solution and 4ul injection was made for MS analysis on TSQ Quantiva using pY LC-MRM/MS method (see above).

### Phosphotyrosine flow through desalt

100 mg (1 cc) Sep-Pak C18 Vac Cartridges (Waters, WAT023590) were conditioned with 1 mL of acetonitrile followed by 1 mL of 0.1% FA / 50% ACN. The cartridges were then equilibrated with four 1 mL injections of 0.1% TFA. pY enrichment flow through samples were acidified with 150 uL of 10% FA and loaded onto the prepared cartridges in two steps of 750 uL. After sample loading, the cartridges were washed three times with 1 mL of 0.1% TFA then one time with 1 mL 1% FA. The samples were then eluted off the cartridges and into 2 mL eppendorf tubes with 2 injections of 750 uL 0.1% FA / 50% ACN. Following elution, the samples were frozen and dried down by vacuum centrifugation. After drying, the samples were reconstituted in 1 mL of 0.1% FA / 3% ACN and the concentration was measured using the Pierce BCA Protein Assay Kit (ThermoFisher, 23225). Using the concentrations measured by the BCA assay, 1 mg aliquots were made for the immobilized metal affinity chromatography (IMAC) enrichment step. The aliquots were frozen and dried down by vacuum centrifugation.

### Immobilized metal affinity chromatography (IMAC) phosphopeptide enrichment

Ni-NTA Agarose beads (Qiagen, 1018244) were gently inverted to resuspend the beads, then 1200 uL slurry, or 600 uL beads, were removed and transferred to a 1.5 mL Eppendorf tube. The beads were spun down for 1 minute at 1500 rcf. The supernatant was removed and the beads were washed three times by adding 1 mL of water onto the beads, inverting the tube to suspend the beads, then spinning down and removing the supernatant. After washing, the beads were stripped of the nickel by incubating end-over-end with 1200 uL of 100 mM ethylenediaminetetraacetic acid (EDTA, Sigma-Aldrich, E7889) at room temperature for 30 minutes. Following the EDTA incubation, the beads were washed three times with HPLC water then incubated with 1200 uL of 10 mM FeCl3 end-over-end at room temperature for 30 minutes. The agarose beads were then washed three times with HPLC water and resuspended with 1:1:1 acetonitrile: methanol: 0.01% acetic acid to a ratio of 1:3 beads to slurry volume. 60 uL slurry, or 20 uL beads, were then aliquotted into 1.5 mL Eppendorf tubes for each 1 mg sample undergoing phosphopeptide enrichment.

The dried peptide aliquots were reconstituted in 0.1% TFA / 50% ACN and vortexed until the peptides were fully dissolved. 0.1% TFA / 100% ACN was then added to each aliquot to bring the final concentration to 0.5 mg/mL in 80% ACN solution. Following reconstitution, 30 fmol of heavy labeled IMAC peptides were spiked into each sample, and the peptide solutions were added on top of the prepared beads and incubated end-over-end for 30 minutes at room temperature. Once the incubation was complete, the beads were spun down for 1 minute at 1500 rcf. The supernatant was removed and saved, and the beads were brought back up in 200 uL of 0.1 % TFA / 80% ACN. The beads were then transferred onto a prepared stagetip for desalting.

### Immobilized metal affinity chromatography stagetip desalt

Stagetips, prepared with two Empore C18 (3M, 2315) punches, were conditioned with 100 uL of methanol and spun down at 3100 rcf for 1 minute. The tips were further conditions with 50 uL of 0.1% FA / 50% ACN, then equilibrated with 2 injections of 100 uL 1% FA. The samples were then loaded onto the stagetips and spun down. The stagetips were desalted with two 50 uL injections of 0.1% TFA / 80% ACN then one 50 uL injection of 1% FA. After desalting, the phosphopeptides were eluted off the agarose beads and onto the stagetips by three 70 uL injections of 500 mM K_2_PO_4_. The samples were washed once with 100 uL of 1% FA, then eluted off the tips with 60 uL 0.1% FA / 50% ACN. The eluates were transferred from Eppendorf tubes to HPLC vials, frozen, and dried down by speedvac. Each sample was then reconstituted in 9 uL of 0.1% FA / 3% ACN solution and 4ul injection was made for MS analysis on TSQ Quantiva using IMAC LC-MRM/MS method (see above).

#### Cell line processing for testing the assay

10 cell lines (PC9, H3122, TMD8, Mino, PC3, OVCAR4, WM266.4, Mel-juso, A375, and RT112) were lysed, digested as described above. 5mg of each was enriched by pY antibody and 1mg of the flow through of that was enriched by IMAC and analyzed following SigPath workflow as described above.

#### Cell line perturbagen sample processing

H3122 and Ls513 cells were treated with either DMSO or drug for 6 hours and 24hours (Figure S 2). H3122 cells were treated with Ceritinib at 300nM concentration and Ls513 cells were treated with Trametinib at 30nM concentration.

Treatments and time points were done in two process replicates. Cells were collected, lysed, digested as described above. Following digestion 5mg of each sample was enriched with pY antibody and 1mg of the flowthrough of that by IMAC and analyzed according to the SigPath workflow described above.

#### Breast cancer xenograft (PDX) tissue processing

Six models were selected from Washington University human to mouse (WHIM) PDX collection (Mundt et al) (4, 30, 21, 6, 2, and 12). Each of the models was treated either with Buperlasib or vehicle. For the 2hr treatment group, the animals received one dose and tissue was collected 2 hours after the treatment either by Buperlasib or vehicle. For the 50hr group, animals received Buperlasib or vehicle at 0hr, 24hr and 48hr, and the tissue was collected at 50 hours. Only in the washout group at 48hr the animals were treated with vehicle instead of the drug. Tissue lysis and digestion was performed as described above. Input peptide amount for pY antibody enrichment varied as follows for the different WHIM models due to limited amount availability of some of the samples: WHIM4 – 4.5mg; WHIM30 – 4.5mg; WHIM21 – 5mg; WHIM6 – 5mg; WHIM2 – 2mg; WHIM12 – 4mg. pY antibody enrichment for the 5 time point and treatment samples Input peptide amount for IMAC was 1mg for all of the samples. pY antibody enrichment for all the five samples of each WHIM were done the same day, but on different days for the different WHIMs. IMAC enrichment was performed over 3 days samples for 2 WHIM models per day.

#### Medulloblastoma tissue processing

39 tissue samples from medulloblastoma patients belonging to all 4 groups (Sonic Hedgehog (SHH), group 3 (GR3), group 4 (GR5) and WNT) were digested as described above. Less than 1mg digested peptides were available for this study and therefore, the pY antibody portion of the procedure was skipped and only IMAC part of the assay was applied to all the samples with 500ug input digested peptides for 85% of them. For 6 of the samples 500ug was not available, therefore the peptide input varied as follows: MB088 (SHH) = 357ug; MB136 (SHH) = 347ug; MB206 (SHH) = 313ug; MB284 (SHH) = 444ug; MB287 (SHH) = 425ug; MB091 (GR4) = 485ug. Sample processing and data acquisition was performed in 4 batches. Samples were randomized in 4 batches making sure to include an equal number of samples from each group.

### Data processing

All analyses of raw mass spectrometry data were performed in Skyline Targeted Mass Spec Environment ((Broudy et al., 2014)). Peak area ratios of endogenous light to stable isotope labeled (SIL) heavy internal standard peptide were calculated in Skyline (Skyline version 4.2.1.18329, https://brendanxuw1.gs.washington.edu/labkey/project/home/software/Skyline/begin.view). All the peaks were manually inspected to make sure accurate and equal integration of light and heavy versions. Peak area ratios of the most abundant, interference free transitions were used for further statistical analysis.

## Statistical analysis

Peak area ratios were log2 transformed and median normalized. A two sample moderated T-test was applied to cell line and PDX datasets using Protigy.

(https://github.com/broadinstitute/protigy). Derived p-values were adjusted for multiple hypothesis testing using Benjamni-Hockberg strategy. Significance was assessed with BHg corrected p-value of less than 0.05.

A one-way ANOVA with an *ad hoc* Tukey’s test (with adjusted p-values for multiple comparisons) was performed in GraphPadPrizm on medulloblastoma dataset. Heatmaps are generated using Morpheus online tool (Morpheus, https://software.broadinstitute.org/morpheus

